# Spatially Aware Adjusted Rand Index for Evaluating Spatial Transcriptomics Clustering

**DOI:** 10.1101/2025.03.25.645156

**Authors:** Yinqiao Yan, Xiangnan Feng, Xiangyu Luo

## Abstract

The spatial transcriptomics (ST) clustering plays a crucial role in elucidating the tissue spatial heterogeneity. An accurate ST clustering result can greatly benefit downstream biological analyses. As various ST clustering approaches are proposed in recent years, comparing their clustering accuracy becomes important in benchmarking studies. However, the widely used metric, adjusted Rand index (ARI), totally ignores the spatial information in ST data, which prevents ARI from fully evaluating spatial ST clustering methods. We propose a spatially aware Rand index (spRI) as well as spARI that incorporate the spatial distance information. Specifically, when comparing two partitions, spRI provides a disagreement object pair with a weight relying on the distance of the two objects, whereas Rand index assigns a zero weight to it. This spatially aware feature of spRI adaptively differentiates disagreement object pairs based on their distinct distances, providing a useful evaluation metric that favors spatial coherence of clustering. The spARI is obtained by adjusting spRI for random chances such that its expectation takes zero under an appropriate null model. Statistical properties of spRI and spARI are discussed. The applications to simulation study and two ST datasets demonstrate the improved utilities of spARI compared to ARI in evaluating ST clustering methods. The R package to compute spRI and spARI is available at https://github.com/yinqiaoyan/spARI.

## 1 Introduction

The spatial transcriptomics (ST) provides unprecedented opportunity to elucidate the spatial context of gene expressions thanks to the advances in biological technologies such as in situ hybridization, in situ sequencing, and spatial barcoding (Williams et al., 2022). Depending on the spatial resolution, ST data can be classified into the spot level and the single-cell level, where in the previous case spots are arranged in a regular lattice format and in the latter case cells distribute irregularly in the tissue section. The ST clustering for spots or cells is an important step in the statistical analysis to understand and investigate the solid tissue spatial heterogeneity especially for cancer and brain. For spot level ST data, the ST clustering becomes region segmentation that aims to partition the tissue section into disjoint domains with different gene expression or morphological features. For single-cell ST data, the ST clustering comes in the form of spatial cell typing that assigns cells to different types. Therefore, downstream biological analyses that rely on the elucidation of spatial heterogeneity, such as marker gene detection (Song et al., 2023), domain/cell type annotation (Shi et al., 2023), imputation (Wan et al., 2023), and region-specific deconvolution (Ma and Zhou, 2022), would greatly benefit from an accurate ST clustering result.

Various ST clustering approaches have been developed in recent several years for region segmentation (Hu et al., 2021; Zhao et al., 2021; Dong and Zhang, 2022; Chidester et al., 2023; Yan and Luo, 2024; Ma and Zhou, 2024) as well as spatial cell typing (Teng et al., 2022; Dong and Zhang, 2022; Li and Zhou, 2022; Singhal et al., 2024; Guo et al., 2024; Yan and Luo, 2025). Different from clustering methods for single-cell RNA-sequencing data, these ST clustering approaches can incorporate spatial information (i.e., the neighbors or spatial coordinates of spots/cells) to improve accuracy, thus offering more insights into the complex disease molecular mechanisms (Park et al., 2023). Considering a large volume of ST clustering methods available, a natural and practical question is that how to compare their performances in a reasonable and quantitative way to provide guidance for an appropriate choice of these ST clustering methods in diverse situations (Yuan et al., 2024).

To evaluate the clustering accuracy, several statistical metrics have been come up with to quantify the comparison of the clustering result against a reference. The Rand index (Rand, 1971) is the first attempt to compare two partitions for a given number of objects, which are spots or cells in our situation, and quantify the “similarity” of the two partitions. Specifically, for two different objects, we first observe whether the two objects are in the same cluster for one partition but are in different groups for the other partition. If it is the case, such two objects are then called a disagreement pair; otherwise, they are named an agreement pair. Subsequently, the Rand index is defined as the ratio of the number of agreement pairs to the total number of pairs, so it takes values between zero and one, and Rand index one indicates that two partitions are identical. The commonly used adjusted Rand index (ARI) (Hubert and Arabie, 1985) is a variant of the Rand index that is corrected for random chance. By counting the common object numbers of two partitions in a contingency table and assuming a generalized hypergeometric distribution for the table entries with fixed column and row sums, ARI has an expectation zero and is bounded above by one. Other clustering evaluation measures include the normalized mutual information and the V-measure (Wu and Wu, 2020). Sometimes, Moran’s *I* is used to measure spatial autocorrelations in a single partition, but it cannot capture the similarity between two partitions. In the current ST clustering literature, ARI is the most widely used clustering evaluation metric.

However, ARI ignores an invaluable characteristic of the ST partition—the spatial locations of objects, so it cannot evaluate the spatial coherence ability of the clustering methods. For example, two different spatial clusterings can result in the same ARI (Sections 4.1 and 4.2). In the analysis of human dorsolateral prefrontal cortex data (Section 5.1), the omission of spatial information by ARI may lead to unsatisfactory evaluation results, where a method with comparable clustering accuracy but weaker spatial coherence receives a higher ARI value. Therefore, there is an urgent need to propose a subtle *spatially aware* clustering metric that can adequately assess a large number of ST clustering algorithms.

We remark that the focus of this paper is not on benchmarking ST clustering algorithms, but on developing a new clustering metric that can directly make use of spatial information to provide more biologically meaningful measurement. In the field of single-cell RNA-seq data clustering, Wu and Wu (2020) begin to improve the Rand index by accounting for the cell type hierarchy. For a disagreement cell pair, its contribution to the weighted Rand index proposed by Wu and Wu (2020) is not identical to zero but relies on the hierarchical relationship between two cells. However, it is not straightforward to incorporate the spatial information of these cells into wRI due to the biological distinction between the cell type hierarchy and the spatial pattern of spots/cells. Recently, Xu et al. (2024) propose a spatial grouping discrepancy metric for ST data, which enables the measurement of spatial distribution discrepancies between predicted and reference labels. Nevertheless, this metric is primarily for anomalous tissue region detection and is not entirely compatible with the general ST clustering accuracy quantification in our task.

In this paper, we present two novel spatially aware clustering metrics based on Rand index and ARI, which are named spRI and spARI, respectively, to quantitatively evaluate the ST clustering performance. Specifically, inspired by the framework of Wu and Wu (2020), when comparing two partitions, spRI provides a disagreement pair of objects with a weight relying on the spatial distance of the two objects, directly employing the available spatial coordinates. In contrast, the classical Rand index assigns an identical zero weight to all disagreement pairs. This spatially aware feature of spRI successfully differentiates disagreement object pairs based on their distinct distances, which cannot be achieved by the Rand index. We adjust spRI to correct for random chances so that its expectation takes on zero value under an appropriate random null model, which results in spARI. We further investigate the statistical properties of spRI and spARI, including the analytical forms between two similar clusterings, the sensitivity to subsampling, hypothesis testings, and computational efficiency. Importantly, in simulation studies, two spatial clustering results with the same ARI can have different spARI values, demonstrating that spARI prefers a more spatially coherent clustering. We also provide an R package for computing spRI and spARI which is user-friendly for statisticians and biologists. The applications to two ST datasets demonstrate the improved utilities of spARI compared to ARI in evaluating ST clustering methods.

## 2 Method

### 2.1 Review of Rand index and adjusted Rand index

We consider *n* objects (spots or cells) with each object *i* (1 ≤ *i* ≤ *n*) having a spatial coordinate *z*_*i*_. We assume there is a reference partition of the *n* objects, denoted by **U** = {*U*_1_, …, *U*_*r*_, …, *U*_*R*_}. In the reference partition **U**, the *n* objects are clustered into disjoint *R* groups, which represent regions for spots and cell types for cells, and *U*_*r*_ is the set of objects that belong to group *r* (1 ≤ *r* ≤ *R*). The reference partition can be obtained from data sources with highly confident manual region or cell type annotations (Wu and Wu, 2020; Yuan et al., 2024) and thus serves as a gold standard to compare ST clustering algorithms. In addition, some clustering method partitions the same *n* objects into *C* clusters, denoted by **V** = {*V*_1_, …, *V*_*c*_, …, *V*_*C*_}, where the cluster number *C* is not necessarily equal to the group number *R* in the reference.

Based on the two partitions **U** and **V**, for any two different objects *i* and *j*, they are named an *agreement pair* if they are grouped in both the reference and the clustering, i.e., *i, j* ∈ *U*_*r*_ and *i, j* ∈ *V*_*c*_ hold for some *r* and *c*, or if they are separated in both the reference and the clustering, i.e., *i, j* ∉ *U*_*r*_ and *i, j* ∉ *V*_*c*_ hold for any *r* and *c*. On the other hand, if two objects *i* and *j* are grouped in one partition but separated in the other partition, this pair is referred to as a *disagreement* pair, following the notations in Hubert and Arabie (1985). For all 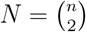 object pairs, according to the “grouped” case or the “separated” case in the reference **U** and the clustering **V**, we have four types for these pairs, and the corresponding contingency table is in Table 1.

**Table 1:** The numbers for the four object pair types. Type “g×g”, the two objects are in the same group in **U** and in the same cluster in **V**. Type “s×s”, the two objects are in different groups in **U** and in different clusters in **V**. Type “s×g”, the two objects are in different groups in **U** but in the same cluster in **V**. Type “g×s”, the two objects are in the same group in **U** but in different clusters in **V**.

| Reference $\mathbf{U}$ \ Clustering $\mathbf{V}$ | | | Sums |
| --- | --- | --- | --- |
|  | Grouped | Separated |  |
| Grouped | “g×g” $N_{11}$ | “g×s” $N_{12}$ | $N_{1\cdot}$ |
| Separated | “s×g” $N_{21}$ | “s×s” $N_{22}$ | $N_{2\cdot}$ |
| Sums | $N_{\cdot 1}$ | $N_{\cdot 2}$ | $N$ |

In Table 1, we denote the object pair numbers of types “g×g”, “s×s”, “s×g” and “g×s” by *N*_11_, *N*_22_, *N*_21_, and *N*_12_, respectively. We also use the dot index to represent the row sums or column sums in Table 1. For example, *N*_1·_ = *N*_11_ + *N*_12_. Subsequently, the number of agreement object pairs is equal to the type “g×g” number plus the type “s×s” number (i.e., *N*_11_ + *N*_22_), while the number of disagreement object pairs becomes the type “s×g” number plus the type “g×s” number (i.e., *N*_21_ + *N*_12_). The Rand index (RI) (Rand, 1971) is then defined as RI = (*N*_11_ + *N*_22_)/*N*, which is exactly the proportion of agreement pairs among all the *N* object pairs.

Rand (1971) and Hubert and Arabie (1985) provide a convenient way to calculate RI using the common object numbers of the two partitions. In Table 2, *n*_*rc*_ is the number of common objects in group *U*_*r*_ and cluster 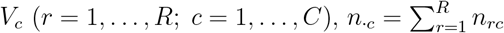, and 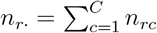. Using these notations, it is easy to derive that

**Table 2:** The common object number table by comparing the reference partition **U** and the clustering partition **V**.

| Reference \ Clustering | | $\mathbf{V}$ | | | | Sums |
| --- | --- | --- | --- | --- | --- | --- |
| | | $V_1$ | $V_2$ | ⋯ | $V_C$ | |
| $\mathbf{U}$ | $U_1$ | $n_{11}$ | $n_{12}$ | ⋯ | $n_{1C}$ | $n_{1\cdot}$ |
| | $U_2$ | $n_{21}$ | $n_{22}$ | ⋯ | $n_{2C}$ | $n_{2\cdot}$ |
| | ⋮ | ⋮ | ⋮ | ⋮ | ⋮ | ⋮ |
| | $U_R$ | $n_{R1}$ | $n_{R2}$ | ⋯ | $n_{RC}$ | $n_{R\cdot}$ |
| Sums | | $n_{\cdot 1}$ | $n_{\cdot 2}$ | ⋯ | $n_{\cdot C}$ | $n_{\cdot\cdot} = n$ |

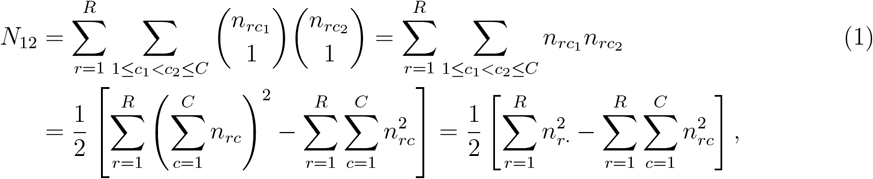

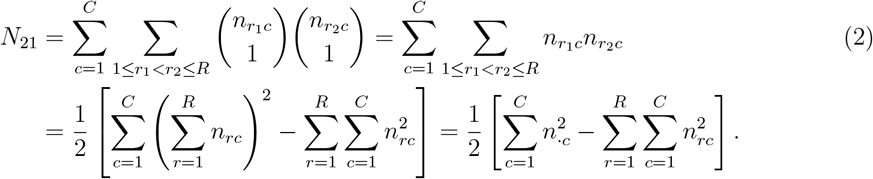

In Equation (1),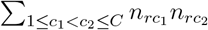 represents the number of object pairs that are in group *U*_*r*_ of **U** but are separated in **V**. Similarly, in Equation (2),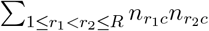 is the number of object pairs that are separated in **U** but are in cluster *V*_*c*_ of **V**. Subsequently,

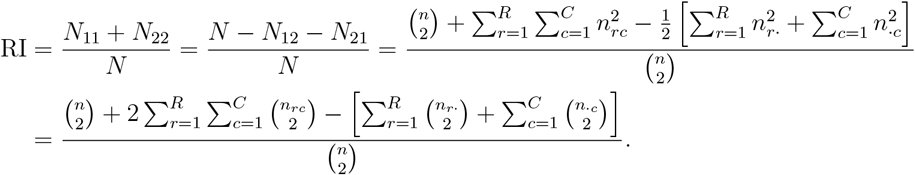

To account for the randomness in the contingency table 2, Hubert and Arabie (1985) set a null model where the common object numbers {*n*_*rc*_ : 1 ≤ *r* ≤ *R*, 1 ≤ *c* ≤ *C*} follow a generalized hypergeometric distribution with the cluster numbers *R* and *C* as well as the column and row sums (*n*_·*c*_ and *n*_*r*·_) fixed. Under this assumption, the marginal distribution of *n*_*rc*_ is actually a hypergeometric distribution, and it turns out that (Hubert and Arabie, 1985) 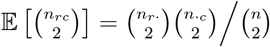. Accordingly, the expectation of RI in the null random model is obtained by

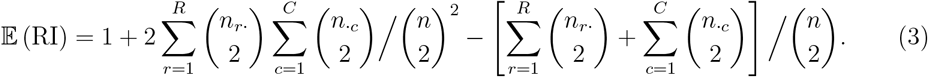

We remark that 𝔼(RI) = 1 if and only if the reference **U** has only one group and all the objects are in one cluster of **V**, i.e., *R* = *C* = 1 and *n*_1·_ = *n*_·1_ = *n*; or each cluster has only one object in both **U** and **V**, i.e., *n*_*r*·_ = *n*_·*c*_ = 1 for ∀*r*, ∀*c*. The proof is in Web Appendix A. When 𝔼(RI) = 1 (two partitions are identical), the adjusted Rand index (ARI) is defined as one; and when E(RI) < 1, ARI has the following form (Hubert and Arabie, 1985),

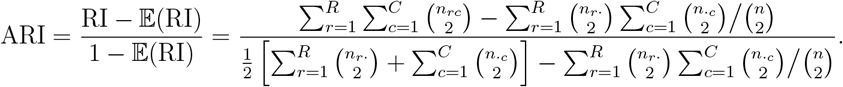

ARI is bounded above by one and has an expectation of zero in the null random model. It is the most widely used clustering evaluation metric in the current ST clustering literature.

### 2.2 The proposed spatially aware adjusted Rand index

In evaluating ST clustering, RI and ARI do not directly account for the spatial information of objects. Following Wu and Wu (2020), we first define a weight function *W* (*i, j*) for each object pair (*i, j*) as follows

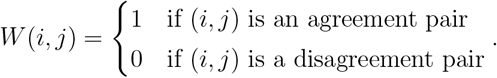

Using this notation, RI can be reformulated as RI = Σ_1≤*i<j*≤*n*_ *W* (*i, j*)/*N*. We observe that for any disagreement pair, its weight in RI is identical to zero, so this weight function cannot make any difference for two disagreement pairs where their spatial distances are distinct. Therefore, we propose a new weight function 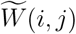 that can incorporate the spatial distance information as follows,

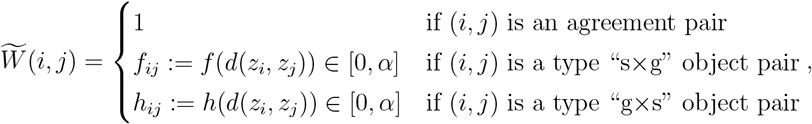

where *d*(*z*_*i*_, *z*_*j*_) is a distance between the spatial locations of object *i* (*z*_*i*_) and object *j* (*z*_*j*_), *f* is a decreasing function from [0, +∞) to [0, α], and *h* is an increasing function from [0, +∞) to [0, α]. The coefficient α belongs to the open interval (0, 1) to keep a positive gap between the maximal weight of the disagreement pair and the weight one of the agreement pair. Throughout this paper, we adopt the Euclidean distance *d*(*z*_*i*_, *z*_*j*_) = ∥*z*_*i*_ − *z*_*j*_∥_2_, *f*(*t*) = α exp{−*t*^2^}, *h*(*t*) = α(1 − exp{−*t*^2^}), and α is fixed at 0.8; the rationales for these choices are provided in the next subsection. To remove the effect of spatial location unit, we normalize *z*_*i*_ by (*z*_*iℓ*_ − min_*i*_ *z*_*iℓ*_)/(max_*i*_ *z*_*iℓ*_ − min_*i*_ *z*_*iℓ*_) for each dimension *ℓ* of *z*_*i*_ such that *z*_*iℓ*_ ∈ [0, 1].

In this definition, even though two disagreement pairs (*i*_1_, *j*_1_) and (*i*_2_, *j*_2_) are of the same disagreement type (“s×g” or “g×s”), their weights are different as long as 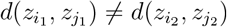. More importantly, the weight function 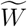 has advantages over *W* in the following two situations which are often encountered in practice.

(i) In situation one where both (*i*_1_, *j*_1_) and (*i*_2_, *j*_2_) are of type “s×g”, if 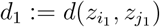 is larger than 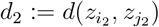, then 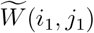 is less than 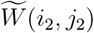 due to the decreasing function *f*. In other words, if two objects are separated in the reference but grouped together in the ST clustering,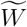 assigns a higher weight to the object pair with a smaller distance. In this “grouped incorrectly” clustering case, we are more willing to forgive an incorrect grouping with a smaller distance.

(ii) In situation two where both (*i*_1_, *j*_1_) and (*i*_2_, *j*_2_) are of type “g×s”, if *d*_1_ > *d*_2_, then 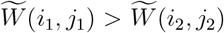 due to the increasing function *h*. In other words, if two objects are in one group in the reference but separated in the ST clustering,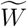 assigns a higher weight to the object pair with a larger distance. In this “separated incorrectly” clustering case, we are more willing to forgive an incorrect separation with a larger distance.

Subsequently, similar to RI, we define the spatially aware Rand index (spRI) as

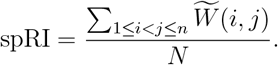

Following the random model used in ARI where column/row sums, the reference group number *R*, and the cluster number *C* in Table 2 are fixed, we aim to calculate the expectation of spRI. We consider two objects (*i, j*) and *i* ≠ *j*. The probability that the two object are in one group of the reference **U** is denoted by *p*_*ij*_, and the probability that they are in one group of the clustering **V** is denoted by *q*_*ij*_. Using this randomness, it turns out that

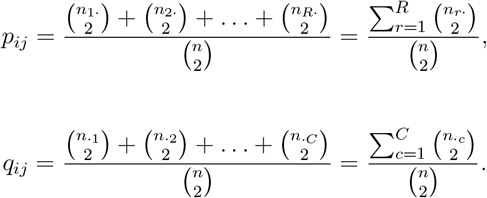

Notice that the terms on the right hand side are not relevant to indices (*i, j*), so we simply let *p*_*ij*_ = *p* and *q*_*ij*_ = *q*. Owing to the independence in assigning a type to an object pair for **U** and **V**, for any object pair, the probability of being type “g×g”, “s×s”, “s×g”, and “g×s” is *pq*, (1 − *p*)(1 − *q*), (1 − *p*)*q*, and *p*(1 − *q*), respectively. Therefore, we have

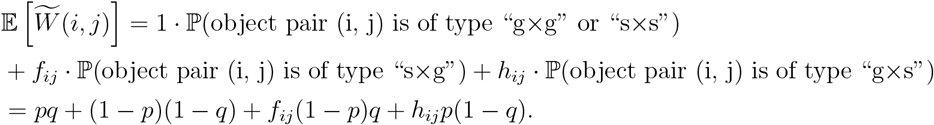

The expectation of spRI is derived as follows,

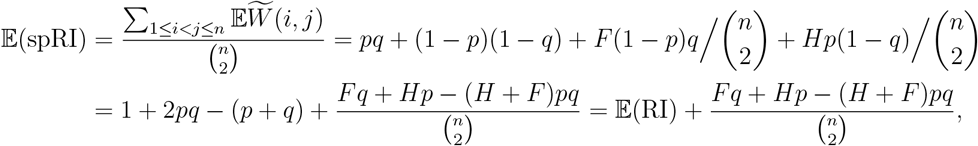

where 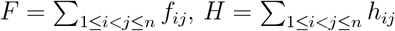 and the last equality holds due to Equation (3). When *f*_*ij*_ = 0 and *h*_*ij*_ = 0 for any (*i, j*), 𝔼(spRI) becomes 𝔼(RI). We notice that for *p, q* ∈ [0, 1], 𝔼(spRI) = 1 if and only if *p* = *q* = 1 or *p* = *q* = 0 (the proof is in Web Appendix B). When 𝔼(spRI) < 1, we define the adjusted version of spRI, named spARI, as

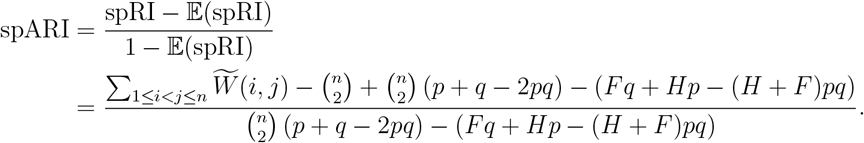

In this way, the expectation of spARI is exactly zero under the null random model, and this property is the same as ARI (Hubert and Arabie, 1985). When 𝔼(spRI) = 1 (two partitions are identical), we define spARI as one. Notice that if we denote the spARI value with **U** in rows and **V** in columns as in Table 2 by spARI(**U, V**), then spARI(**U, V**) ≠ spARI(**V, U**) as “s × g” pairs and “g × s” pairs are not symmetric. Therefore, to ensure uniformity, we always put the reference partition in the first position of spARI.

In practice, when we compare two clustering methods (against the same reference), it is helpful to investigate whether one method with a higher spRI than the other method can still give a higher spARI. Web Appendix C provides a practical guideline about when spRI and spARI can reach the same evaluation conclusion for two ST clustering methods.

**Figure S1:**
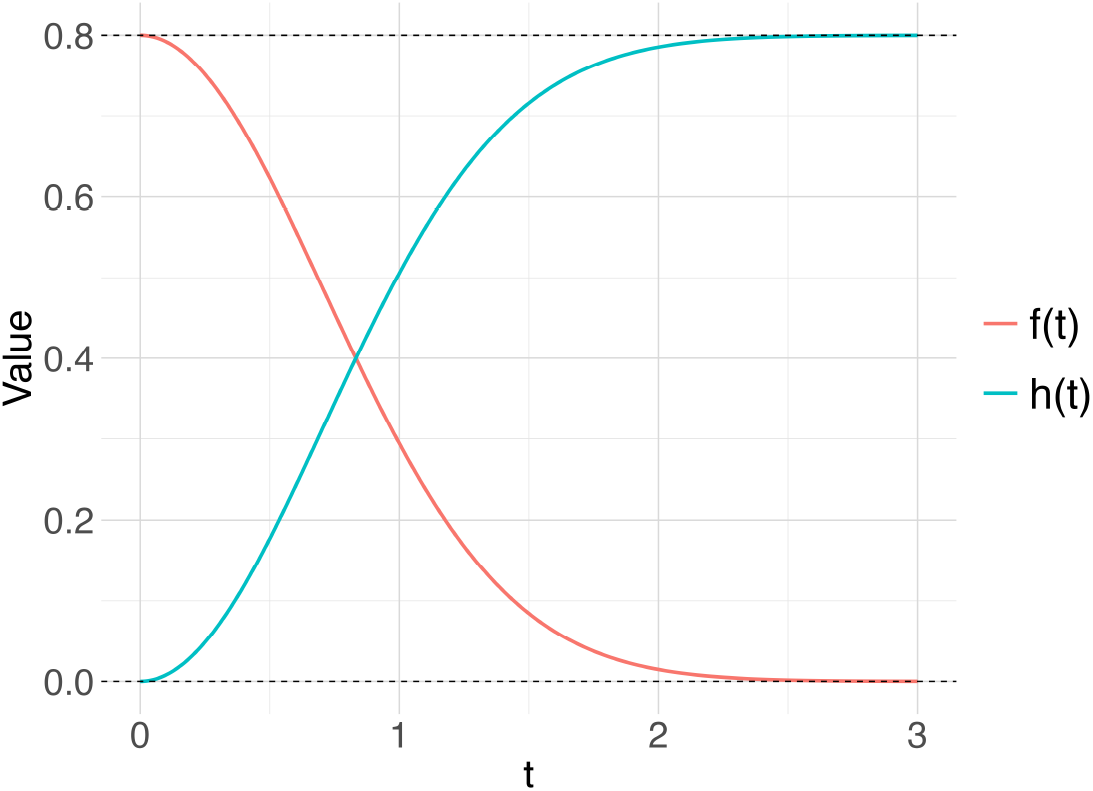
The function curves of *f*(*t*) = α exp{−*t*^2^} and *h*(*t*) = α(1 − exp{−*t*^2^}).

### 2.3 The choices for the parameters in spARI

Currently, most ST datasets are collected from tissue slices in a two-dimensional plane, so the Euclidean distance is a natural choice to compute the pairwise distances between objects when calculating spARI. In general, alternative distances can be employed. For example, the use of Euclidean distance implicitly assumes that all coordinate dimensions share equal weights. As a generalization, the Mahalanobis distance is able to assign different weights or even introduce correlation structures for dimensions through the covariance matrix 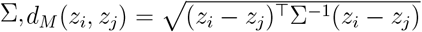. When Σ is a diagonal matrix, the inverse of each diagonal entry serves as a specific weight for the corresponding dimension. In addition, for geospatial data with large spatial extent on the sphere, the geodesic distance may provide a more appropriate measure of spatial proximity than Euclidean distances.

The functions *f* and *h* are defined as *f*(*t*) = α exp{−*t*^2^} and *h*(*t*) = α(1 − exp{−*t*^2^}). Their corresponding function curves are illustrated in Supplementary Figure S1. Both *f* and *h* take values in the interval [0, α], with *f* being decreasing and *h* increasing. The choice of the exponential function exp{−*t*^2^} stems from its rapid decay as *t* increases, which allows spARI to better distinguish between nearby and distant object pairs. Generally, such exponential (or Gaussian) functions are widely employed in spatial statistics to model the decay rate of the spatial correlation with distance (Williams and Rasmussen, 2006). To address the situation where other types of functions may be more appropriate, the R package of spARI now supports user-specified functions *f* and *h* with the following constraints—they should map from their domains to [0, α] with *f* decreasing and *h* increasing.

Regarding the parameter α, we conducted a sensitivity analysis in Web Appendix D to assess its impact on the spARI value. When α approaches zero, the contribution of disagreement pairs to spARI becomes negligible, so spARI becomes close to ARI. In contrast, as α increases toward one, the accumulated weights of disagreement pairs become larger. Thus, we set the default value of α to 0.8 to ensure a reasonable positive gap between the weights of disagreement and agreement pairs, also capturing meaningful spatial information.

In practice, we should be cautious about the alternative choices of the parameters in spARI unless users have strong and reliable domain expertise that can accurately specify the distance function or associated parameters. The introduction of potentially unjustified choices may mislead the evaluation. Since spARI is designed to serve as an objective evaluator of spatial clustering methods, we recommend that the need for manual or data-driven tuning should be minimized as much as possible if we do not have strong prior information.

## 3 Some results for spRI and spARI

### 3.1 Expressions for spRI between two similar clusterings

As all clusterings can be formed by applying a series of operations to an initial clustering with a simple structure, it is important to derive the form of spRI for cases where one clustering is constructed from another based on some basic operation. Following the evaluation framework in Rand (1971), in Web Appendix E, we provide the analytical forms of spRI (a) between the initial clustering and a new clustering formed by simple operations; (b) between two clusterings, each of which is created by applying the same operation to different parts of the same initial clustering; and (c) in some standard settings including all-objects-in-one-cluster, one-object-in-one-cluster, as well as the opposite clustering.

### 3.2 The sensitivity of spRI to subsampling

The value of spRI may change when we focus on a subset of all objects, and this subsampling technique is often adopted in a pilot study when we aim to cluster hundreds of thousands of spots or cells. Hence, it is useful to look into the sensitivity of spRI, which measures how the spRI varies with the subsampling proportion *τ* (0 < *τ* < 1). Let Ω represent the set of all *n* objects, and *τ* Ω the subset of Ω with *nτ* objects. In Table 2, if we use the subsample set *τ* Ω, the expected object number falling into the *r*th row and *c*th column is *τn*_*rc*_. For the ease of computation, we assume all these *τn*_*rc*_ (1 ≤ *r* ≤ *R*, 1 ≤ *c* ≤ *C*) are integers. In Web Appendix F, we prove that under mild conditions the difference between spRI on the subset and spRI on the whole dataset can be small as long as *n* is large enough, and the subsampling approximation error is given by 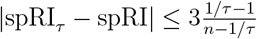.

### 3.3 Hypothesis testings for spARI

In most ST clustering literature, only a single clustering index is calculated and reported for method evaluations. It is useful to investigate whether the clustering partition is significantly associated with the reference partition. This corresponds to testing the null hypothesis that the spARI value is identical to zero (no association between reference and clustering). We denote the spARI value using the observed partition by spARI_obs_, and we test the null hypothesis *H*_0_ : spARI = 0 versus the alternative hypothesis *H*_*a*_ : spARI > 0. Without any assumption about the probability distribution of spARI, we adopt the permutation test (Pitman, 1937) to implement the nonparametric test. The detailed steps in the algorithm are presented in Web Appendix G. Notice that as far as we know, there still does not exist any statistical method that can test the difference between two ARI values (i.e., *H*_0_: ARI_1_=ARI_2_ v.s. *H*_*a*_: ARI_1_ > ARI_2_); developing a powerful hypothesis testing for comparing two spARI values is challenging and left for future work.

### 3.4 Computational efficiency of spARI

The R package “spARI” is developed to encapsulate the computational procedure of spRI and spARI, and is publicly available at https://github.com/yinqiaoyan/spA*R*I. Users only need to input the two partition results along with spatial coordinates or a precomputed distance matrix, and then the main function efficiently computes the spRI and spARI values. The computational complexity of computing ARI is 𝒪 (*n*^2^). In Web Appendix H, we find that if we use a precomputed distance matrix as input, the runtime is under one second for the object number *n* ≤ 8, 000 on a MacBook Air powered by Apple M4 CPU with 16GB of RAM. When *n* = 100, 000, if a sparse distance matrix is passed to the R function to compute the spARI value, the runtime is around 4.8 seconds. This shows that spARI is scalable to large-scale ST data if users are able to provide the distance matrix in a sparse format.

## 4 Simulation study

In this section, we generate three synthetic ST data to illustrate the distinction between RI/ARI and spRI/spARI when evaluating two clustering results. We demonstrate that the proposed spRI/spARI can effectively distinguish structural differences between two clustering partitions using the spatial information of disagreement object pairs, even when their RI/ARI values are identical. For presentation convenience, we denote the sum of weights contributed by type “s×g” object pairs, 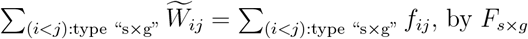, by *F*_*s*×*g*_, and the sum of weights by type “g×s” object pairs,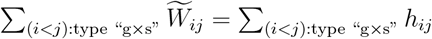, by *H*_*g*×*s*_.

### 4.1 Two clustering results with different *F*_*s*×*g*_ and identical *H*_*g*×*s*_

The reference dataset consists of two groups, each with 6 objects in the left panel of Figure 1(a). Objects numbered 1 to 6 belong to group R1, and the other six objects numbered 7 to 12 belong to group R2. There exist two clustering partitions that are illustrated in the middle and right panels of Figure 1(a), respectively. In clustering result i, object number 5, which is in reference group R1, is incorrectly clustered into C2, while in clustering result ii, object number 2 is incorrectly clustered into C2. Both the clustering results contain only one misclustered object, and they have the same object numbers in C1 and C2, respectively, so their RI and ARI values are identical.

However, the positions of the misclustered objects are different between the two results, leading to distinct spRI and spARI shown in Figure 1(a). Specifically, in clustering result i, the type “s×g” object pairs consist of (5,7), (5,8), (5,9), (5,10), (5,11), and (5,12), giving *F*_*s*×*g*_ = 3.143; and the type “g×s” object pairs include (1,5), (2,5), (3,5), (4,5), and (5, 6), giving *H*_*g*×*s*_ = 0.923. In contrast, in clustering result ii, the type “s×g” object pairs, which are (2,7), (2,8), (2,9), (2,10), (2,11), and (2,12), result in a smaller value of *F*_*s*×*g*_ = 2.065 than that of clustering i because object number 2 is more far away from other objects in cluster C2 compared to object number 5. Type “g×s” object pairs in clustering ii are (1,2), (2,3), (2,4), (2,5), and (2,6), providing an identical *H*_*g*×*s*_ = 0.923 to that in clustering i due to the location symmetry of object number 2 and object number 5 in reference group R1.

**Figure 1.**
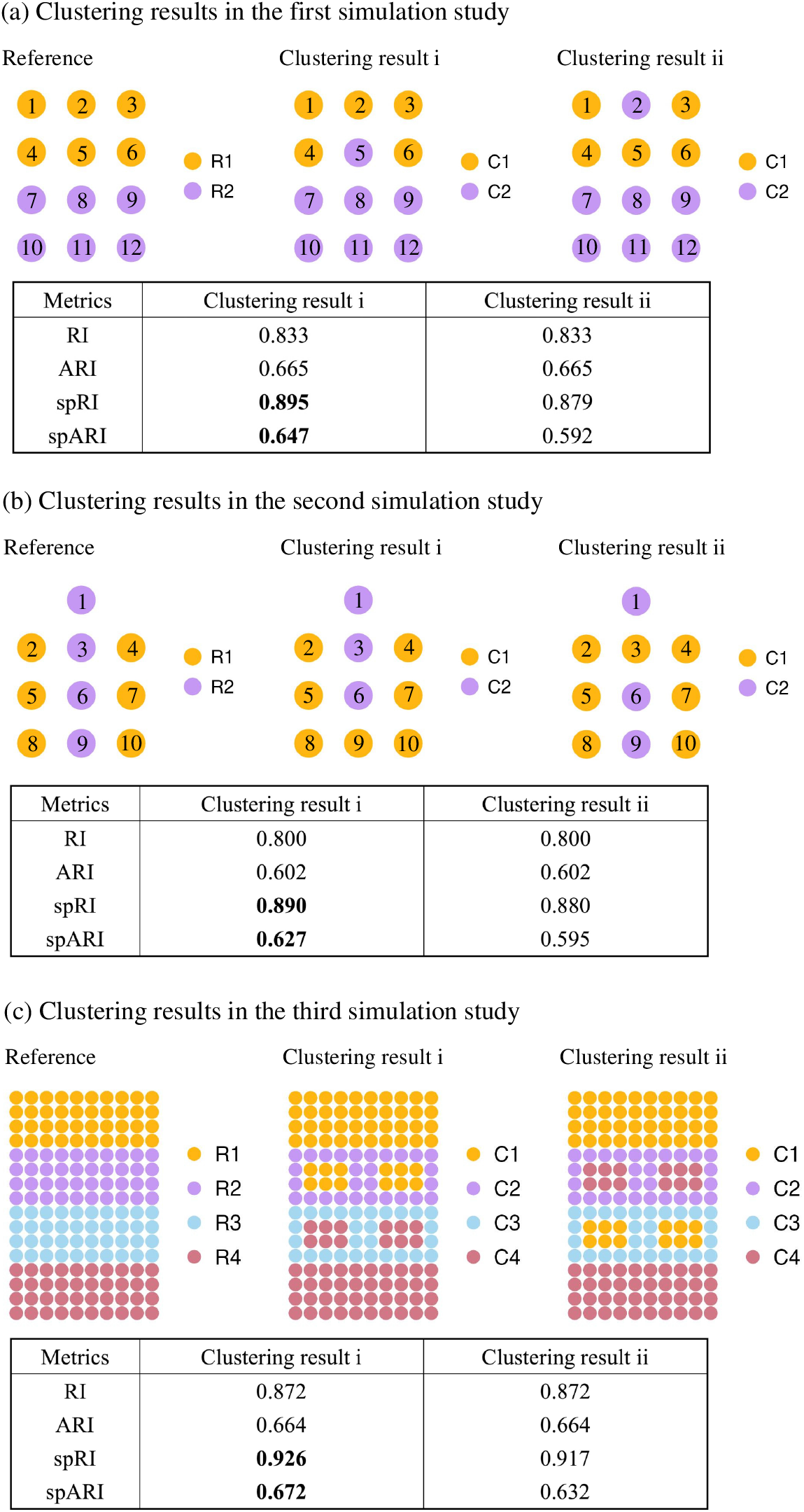
The clustering results in (a) the first, (b) the second, and (c) the third simulation study. There are 12, 10, and 160 objects in the simulated data, respectively. The left, middle, and right panels correspond to the reference partitions of the objects, the clustering partitions by method i, and the clustering partitions by method ii, respectively. The tables at the bottom of panels demonstrate the specific values of RI, ARI, spRI, and spARI for the two clustering results.

Based on the observations above, it is easy to show that the spRI value of clustering result i (0.895) is higher than that of clustering result ii (0.879). Since the expectations of spRI are the same for both the clustering results (i.e., η = 0 in Supplementary Proposition S1), clustering result i also achieves a higher spARI value.

### 4.2 Two clustering results with identical F_*s*×*g*_ and different *H*_*g*×*s*_

In this case, we consider a total of 10 objects. In the reference partition in Figure 1(b), objects numbered 2, 4, 5, 7, 8, and 10 belong to group R1, while objects numbered 1, 3, 6, and 9 in the middle part belong to R2. Similar to the previous case, each of the two clustering results has only one object that is clustered incorrectly in the middle and right panels of Figure 1(b), and they share the same object numbers in C1 and C2, respectively, resulting in their identical RI and ARI values.

In clustering result i, the type “s×g” object pairs include (2,9), (4,9), (5,9), (7,9), (8,9), and (9,10), with a corresponding sum of weights *F*_*s*×*g*_ = 3.160. The type “g×s” object pair comprises (1,9), (3,9), and (6,9), offering a sum of weights *H*_*g*×*s*_ = 0.877. In the meanwhile, in clustering result ii, the type “s×g” object pairs (2,3), (3,4), (3,5), (3,7), (3,8), and (3,10) have a sum of weights, *F*_*s*×*g*_ = 3.160, the same as that in clustering result i, and the type “g×s” object pairs (1,3), (3,6), and (3,9) obtain a sum of weights, *H*_*g*×*s*_ = 0.455, smaller than the counterpart in clustering result i. Thus, clustering result i has a higher spRI value than clustering result ii. Due to the equal expectations of spRI in both clustering results, clustering result i also exhibits a higher spARI value, according to Supplementary Proposition S1.

These two cases demonstrate that even when ARI values are the same, spARI can still effectively evaluate and distinguish various clustering partitions, also highlighting that spARI favors partitions with compact and continuous region structures.

### 4.3 A general clustering setting

We expand the object number to 160 and consider a general setting where there are four reference groups, which mimics the layered structures observed in mouse brain tissue sections; see Figure 1(c). Each layer contains 40 objects. Two clustering results represent the partitions obtained by two ST clustering methods, as shown in the middle and right panels of Figures 1(c), respectively. Each of the two partitions contains 24 incorrectly clustered objects, and the number of objects in each cluster is identical for the two partitions, leading to equal values of RI and ARI. However, in clustering result i, these misclustered objects are spatially closer to objects in the same clusters than those in clustering result ii, so clustering result i has a larger *F*_*s*×*g*_ value. On the other hand, since the distances between misclustered objects and other objects that belong to their reference group remain the same in both clustering partitions, the two clustering results have the same *H*_*g*×*s*_ value. Therefore, the clustering result i achieves higher spRI and spARI values than clustering result ii, suggesting that clustering method i shows more efforts to keep regions integral than clustering method ii.

## 5 Real application

We apply six region segmentation methods—STAGATE (Dong and Zhang, 2022), SpaGCN (Hu et al., 2021), BayesSpace (Zhao et al., 2021), BANKSY (Singhal et al., 2024), BINRES (Yan and Luo, 2024), and IRIS (Ma and Zhou, 2024)—to a spot level ST dataset, and five cell typing methods—STAGATE, SpaGCN, BANKSY, BACT (Yan and Luo, 2025), and BASS (Li and Zhou, 2022)—to a single-cell level ST dataset, and their ARI and spARI values are then calculated and compared. We remark that this application does not perform a benchmarking study for existing ST clustering but demonstrates the advantages of spARI in capturing spatial information of objects over ARI.

### 5.1 Region segmentation evaluation in human dorsolateral prefrontal cortex data

The human dorsolateral prefrontal cortex (DLPFC) is a functional area of the human brain involved in cognitive control and attention regulation. The dataset contains twelve DLPFC tissue sections with manual annotations (reference) provided by Maynard et al. (2021). The DLPFC section 151509 is used for downstream analysis, which contains 4789 spots and 33538 genes. The upper panel of Figure 2(a) shows the bar plots of the ARI and spARI values for each method. Based on ARI, the method performance ranking in this dataset is IRIS, SpaGCN, STAGATE, BINRES, BANKSY, and BayesSpace, while the ranking becomes IRIS, STAGATE, SpaGCN, BINRES, BayesSpace, and BANKSY using spARI. Specifically, BANKSY has a higher ARI value (0.357) than BayesSpace (0.343), but its spARI (0.342) is lower than BayesSpace (0.359).

**Figure 2.**
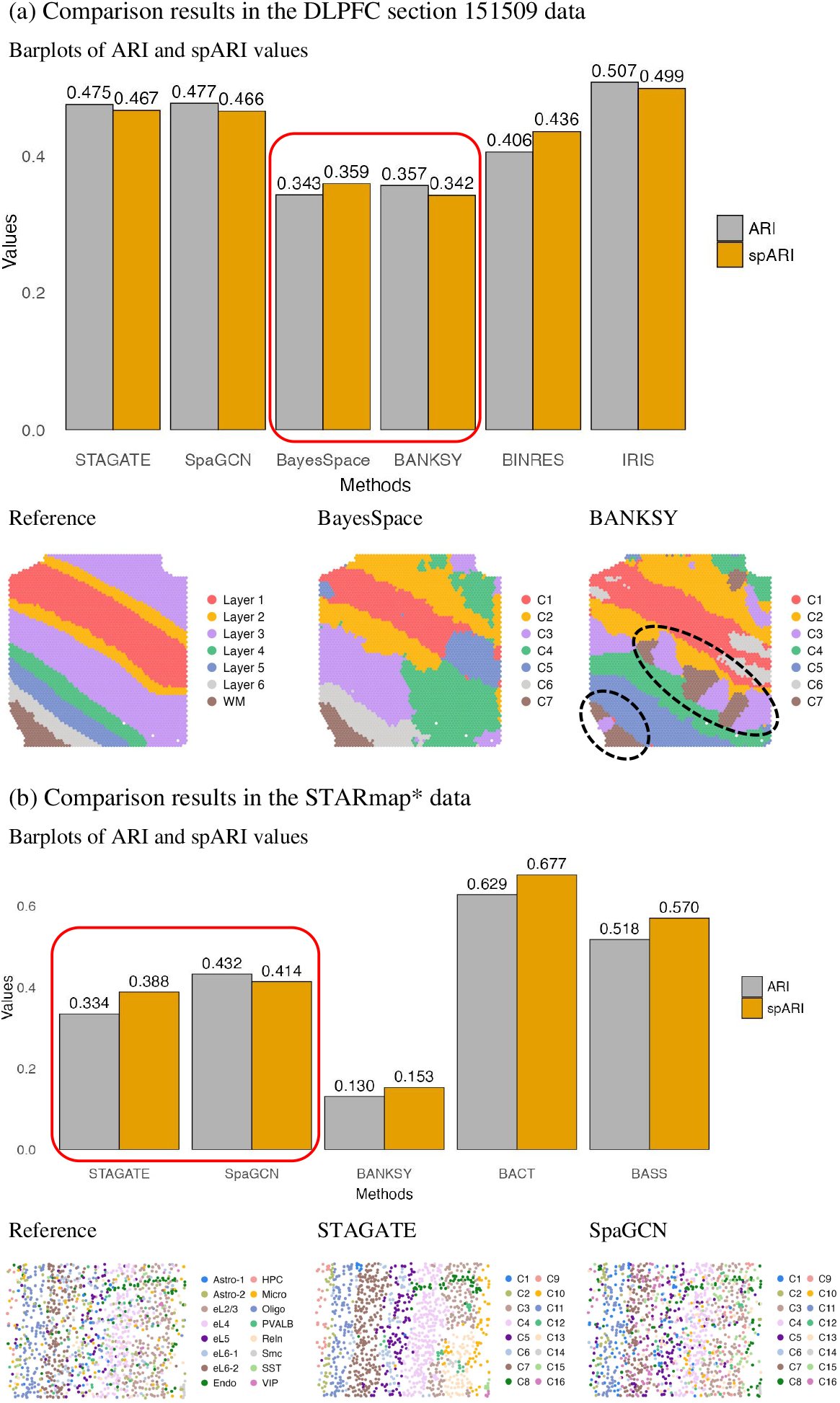
(a) The comparison results in the DLPFC section 151509 data. The upper panel displays the barplots of ARI and spARI values for the six region segmentation methods— STAGATE, SpaGCN, BayesSpace, BANKSY, BINRES, and IRIS. The ARI/spARI values are displayed at the top of bars. The lower panels from left to right show the reference annotations of the DLPFC 151509 section (including six DLPFC layers and the white matter (WM)), the region segmentation by BayesSpace, and the region segmentation by BANKSY. (b) The comparison results in the STARmap* data. The upper panel shows the barplots of ARI and spARI values for the five cell typing methods—STAGATE, SpaGCN, BANKSY, BACT, and BASS. The ARI/spARI values are displayed at the top of bars. The lower panels from left to right show the reference annotations of the STARmap* data (including sixteen cell types), the cell typing by STAGATE, and the cell typing by SpaGCN.

To clarify the discrepancy between ARI and spARI for BayesSpace and BANKSY, we visualize their spatial partitions in the lower middle and right panels of Figure 2(a) to provide further insights. We also show the number of agreement pairs denoted by #{s×s, g×g}, the sum of weights f_*ij*_ for type “s×g” object pairs (*F*_*s*×*g*_), the sum of weights *h*_*ij*_ for type “g×s” object pairs (*H*_*g*×*s*_), spRI, 𝔼(spRI), and spARI in the first two rows of Table 3.

**Table 3:** Related quantities used to calculate the spARI for BayesSpace and BANKSY in the analysis of DLPFC section 151509 data and for STAGATE and SpaGCN in the analysis of STARmap* data.

| Data | Methods | $\#\{s \times s, g \times g\} / \binom{n}{2}$ | $H_{g \times s} / \binom{n}{2}$ | $F_{s \times g} / \binom{n}{2}$ | spRI | $\mathbb{E}(\text{spRI})$ | spARI |
| --- | --- | --- | --- | --- | --- | --- | --- |
| DLPFC | BayesSpace | 0.7760 | **0.0301** | **0.0550** | **0.8611** | 0.7832 | **0.3594** |
|  | BANKSY | **0.7874** | 0.0294 | 0.0420 | 0.8588 | **0.7853** | 0.3421 |
| STARmap* | STAGATE | 0.8684 | **0.0103** | **0.0509** | 0.9296 | 0.8849 | 0.3884 |
|  | SpaGCN | **0.9037** | 0.0094 | 0.0251 | **0.9382** | **0.8947** | **0.4135** |

Based on the reference illustrated in the lower left panel of Figure 2(a), we observe that both BayesSpace and BANKSY identify the major part of Layer 1, but exhibit unsatisfactory performances in detecting Layers 2 and 3. For Layers 4, 5 and 6, neither method captures the complete three-layer structure well. Quantitatively, the first two rows of Table 3 show that BANKSY has a larger number of agreement pairs than BayesSpace. Nevertheless, the dashed domains in the lower right panel of Figure 2(a) tell us that regions C3 and C7 in the BANKSY result are spatially fragmented. This fragmentation leads to a smaller value of *H*_*g*×*s*_ for the type “g×s” object pairs and also a smaller value of *F*_*s*×*g*_ for the type “s×g” object pairs than BayesSpace. In contrast, the region structures in BayesSpace’s partitions look more integral. Consequently, although BayesSpace has a lower ARI, BayesSpace can attain a larger spARI score (0.3593) than BANKSY (0.3420) by effectively incorporating the spatial information. These findings are further supported by the biological interpretations provided in Web Appendix I. This indicates that spARI prefers clustering methods that can yield more compact spatial structures.

### 5.2 Cell typing evaluation in mouse visual cortex STARmap* data

STARmap* is an imaging-based ST technique that captures gene expression patterns of single cells (Wang et al., 2018). This dataset consists of 1207 cells and 1020 genes. The ARI/spARI barplots of cell typing methods are demonstrated in the upper panel of Figure 2(a). The ranking of these methods using ARI is the same as that using spARI, meaning that the evaluation results by ARI and spARI are consistent in this case. This also indicates that the incorporation of spatial information into ARI does not change the ranking of these cell typing methods. However, there are still some interesting observations. For example, when comparing SpaGCN and STAGATE, their ARI difference is 0.098, but the difference decreases to 0.026 using spARI.

To look into what makes this difference, we provide the reference and the clustering results by STAGATE and SpaGCN in the lower panels of Figure 2(b) as well as related quantities during the calculations for their spARI. We can observe that in the lower middle and right panels of Figure 2(b), the cell cluster boundaries in STAGATE (e.g., clusters C3 and C4) are more clear than those in SpaGCN. In other words, more spatially coherent regions are formed in STAGATE. Nevertheless, as spRI is a trade off between RI and spatial proximity, the spRI of STAGATE is still lower than that of SpaGCN (last two rows of Table 3). To put it another way, in the lower left panel of Figure 2(b), some reference cell types exhibit irregular and scattered distributions in the tissue section (e.g., Astro-2, Micro, and Endo) rather than forming well-partitioned layers. Consequently, the more compact clustering characteristic of STAGATE does not enable it to surpass SpaGCN in terms of spRI scores.

Moreover, according to Supplementary Proposition S1, SpaGCN has a larger spARI value than STAGATE. Although the ranking of STAGATE and SpaGCN remains unchanged for ARI and spARI, the narrowed gap in spARI values compared to ARI indicates that spARI not only assesses the clustering accuracy but also captures the structural information of the resulting spatial clusters, demonstrating its utility for a thorough evaluation of the ST clustering.

## 6 Discussion

ARI has been widely used for comparing clustering methods, but it lacks spatial awareness, which prevents it from fully evaluating and comparing spatial clustering methods. In this paper, we propose the spatially aware adjusted Rand index, spARI, to evaluate the ST clustering methods. Built upon ARI that only uses the partitions to provide a clustering accuracy index, spARI can further incorporate the spatial information to assess the ST clustering accuracy. Specifically, the weights in spARI for disagreement object pairs depend on the the distances of objects instead of being identically zero in ARI. It is this feature that equips spARI with the ability to differentiate two ST clustering methods with the same ARI value and that makes spARI in favor of a more spatially coherent clustering. Related properties for spRI and spARI are provided and discussed. Simulation studies and real applications demonstrate that spARI not only considers the consistency of two partitions as reflected by ARI but also takes care of the ability to preserve continuous spatial structures. More importantly, the application of spARI is not restricted to the spatial transcriptomics field but actually can be adopted in any spatial clustering area, such as meteorological data clustering (Pan et al., 2024) and geographical data clustering (Kang et al., 2022) (see Web Appendix J). Therefore, we envision that spARI will be an effective and statistically sound index for any problems associated with spatial clustering evaluations.

## Acknowledgements

We thank Editor, Associate Editor, and reviewers for their insightful suggestions that have significantly improved the quality of this paper. This research was supported by National Natural Science Foundation of China with grant numbers 12471275 and 72271060.

## Supplementary Materials

Web Appendices, Figures, code and data referenced in the paper are available. The R package to compute spRI and spARI is available at https://github.com/yinqiaoyan/spA*R*I.

## Data Availability

All data used in this paper are available in the supplementary materials.

## Supplementary Materials

## Web Appendix A: Proof of the sufficient and necessary conditions for 𝔼(RI) = 1

*Proof*. We first define that 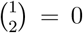 and observe that *n*_*r*·_ ≥ 1 and *n*_·*c*_ ≥ 1. By definition, 𝔼(RI) = 1 if and only if

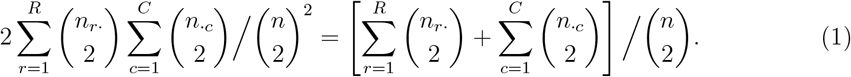

Multiplying both sides of Equation (1) by 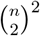 yields

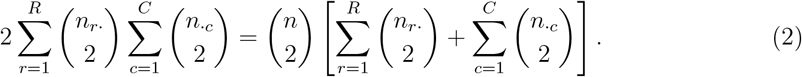

Next, we examine whether this equation holds under the following four distinct cases.

Case 1. *R* = 1 and *C* = 1. In this case, *n*_*r*·_ = *n*_·*c*_ = *n*, and hence Equation (2) holds directly.

Case 2. *R* = 1 and *C* > 1. Here, *n*_*r*·_ = *n*. Equation (2) then implies that 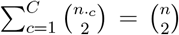.However, it is straightforward to verify that the inequality 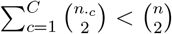 holds whenever *C* > 1, which leads to a contradiction. Thus, Equation (2) does not hold in this case.

Case 3. *R* > 1 and *C* = 1. Similar to Case 2, we obtain that *n*_·*c*_ = *n* and Equation (2) implies that 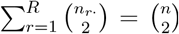, which again contradicts the inequality 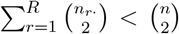 when *R* > 1. Therefore, Equation (2) does not hold in this case either.

Case 4. *R* > 1 and *C* > 1. Let 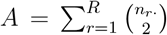 and 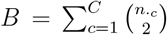, both nonnegative and strictly less than 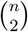. Then the following two inequalities hold: 2 min(*A, B*) ≤ *A* + *B* and 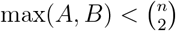, which yields

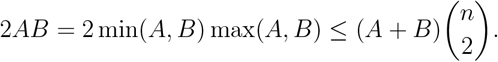

The equality holds (i.e., Equation (2)) if and only if *A* = 0 and *B* = 0, which are satisfied only when *n*_*r*·_ = 1 for all 1 ≤ *r* ≤ *R* and *n*_·*c*_ = 1 for all 1 ≤ *c* ≤ *C*.

Therefore, we conclude that 𝔼(RI) = 1 if and only if one of the following two conditions holds: (i) the reference **U** has only one group and all the objects are in one cluster of **V**, i.e., *R* = *C* = 1 and *n*_1·_ = *n*_·1_ = *n*; or (ii) each cluster has only one object in both **U** and **V**, i.e., *R* = *C* = *n* and *n*_*r*·_ = *n*_·*c*_ = 1 for all 1 ≤ *r* ≤ *R* and 1 ≤ *c* ≤ *C*.

## Web Appendix B: Proof of the sufficient and necessary conditions for 𝔼(spRI) = 1

*Proof*. We recall that the expectation of spRI is formulated as:

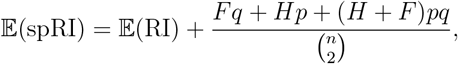

where 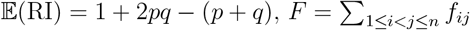 and 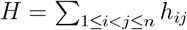. Then 𝔼 (spRI) = 1 holds if and only if

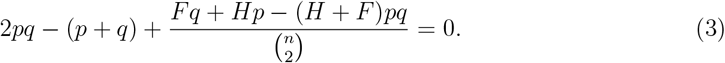

We consider the following four cases to identify whether this equality holds.

Case 1. *R* = 1 and *C* = 1. In this setting, *n*_*r*·_ = *n*_·*c*_ = *n* and *p* = *q* = 1. We then obtain that Equation (3) holds directly.

Case 2. *R* = 1 and *C* > 1. We have *n*_*r*·_ = *n, p* = 1 and *q* ∈ [0, 1). Substituting these values into Equation (3) gives 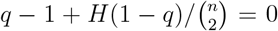, which implies that 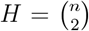. This contradicts the fact that 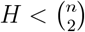 when *C* > 1, so Equation (3) does not hold in this case.

Case 3. *R* > 1 and *C* = 1. We similarly obtain that *n*_·*c*_ = *n, q* = 1 and *p* ∈ [0, 1), and Equation (3) implies that 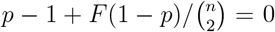 which again leads to a contradiction, since 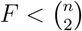 when *R* > 1. Thus, Equation (3) does not hold in this case either.

Case 4. *R* > 1 and *C* > 1. In this case, *p, q* ∈ [0, 1). Through Equation (3), *q* can be expressed as a function of *p*

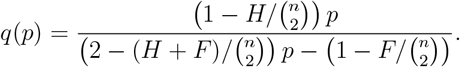

To ensure that the denominator is nonzero, *p* must not equal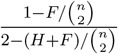, which is denoted by *p*^∗^.Given that 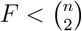 and 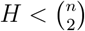, it follows that 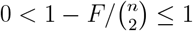 and 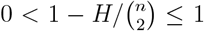, and it is easy to verify that 0 < *p*^∗^ < 1. The derivative of *q*(*p*) is given by

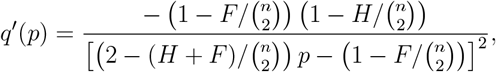

which is negative in the domain of *q*(*p*), indicating that *q* is decreasing on each of the intervals [0, *p*^∗^) and (*p*^∗^, 1) with respective to *p*.

Moreover, from the form of *q*(*p*), we observe that lim_*p*→1−_ *q*(*p*) = 1 and lim_*p*→0+_ *q*(*p*) = 0. Since both *p* and *q* are restricted to the interval [0, 1), the only point in this domain that satisfies the expression of *q*(*p*) is *p* = *q* = 0, under which Equation (3) is satisfied.

Thus, 𝔼(spRI) = 1 holds if and only if one of the following two conditions holds: (i) the reference **U** consists of a single group and all objects in **V** belong to one cluster, i.e., *R* = *C* = 1 and *n*_1·_ = *n*_·1_ = *n* (*p* = *q* = 1); or (ii) every cluster in both **U** and **V** contains only one object, i.e., *R* = *C* = *n* and *n*_*r*·_ = *n*_·*c*_ = 1 for all 1 ≤ *r* ≤ *R* and 1 ≤ *c* ≤ *C* (*p* = *q* = 0).

## Web Appendix C: Consistency between spRI and spARI in method comparison

We aim to determine whether a method with a higher spRI can consistently yield a higher spARI compared to another method. The following proposition answers the question.

### Proposition S1.

*Consider two clustering partitions* **V**_1_ *and* **V**_2_. *We denote the spRI and spARI values of* **V**_1_ *by spRI*_1_ *and spARI*_1_, *respectively, and those of* **V**_2_ *by spRI*_2_ *and spARI*_2_, *respectively. Let* ξ = *spRI*_1_ −*spRI*_2_ *and* η = 𝔼(*spRI*_1_) − 𝔼(*spRI*_2_), *then spARI*_1_ > *spARI*_2_ *if and only if* 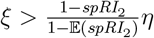.

*Proof*. Using the notations ξ and η, we can express spARI_1_ in the following form

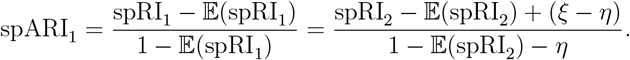

Accordingly, spARI_1_ > spARI_2_ if and only if

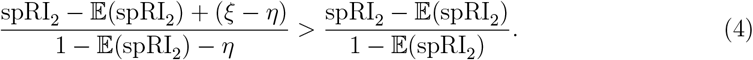

As mentioned in the main text, unless in the cases *R* = *C* = 1 or *R* = *C* = *n*, the expected value of spRI is strictly less than one, implying that 1 − 𝔼(spRI_1_) and 1 − 𝔼(spRI_2_) are strictly positive. Next, we multiply both sides in inequality (4) by (1 − 𝔼(spRI_1_))(1 − 𝔼(spRI_2_)), resulting in

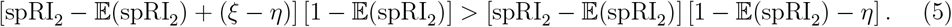

After simple algebra and using the positivity of 1 − 𝔼(spRI_2_), we can obtain that the inequality (5) holds if and only if

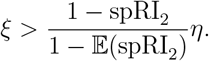

□

Without loss of generality, we assume that the clustering partition **V**_1_ has a higher spRI value than **V**_2_, i.e., ξ > 0. Based on Proposition S1, when η ≤ 0, it is evident that **V**_1_ also gives a higher spARI value. When η > 0, **V**_1_ has a higher spARI value if 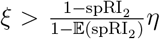 is satisfied. This provides a practical guideline about when spRI and spARI can reach the same evaluation conclusion for two ST clustering methods.

## Web Appendix D: Sensitivity analysis for α

Regarding the parameter α, we conducted a sensitivity analysis based on the third simulation study (Section 4.3 in the main text) to assess its impact on the spARI value. Specifically, we varied α over the set {0.2, 0.3, 0.4, 0.5, 0.6, 0.7, 0.8, 0.9} and computed the spARI value of the first clustering partition for each α. We also calculated the ARI value of the first clustering result, which does not change with α. Figure S2 illustrates the difference between spARI and ARI, and this difference becomes more noticeable as α increases. This phenomenon arises because α controls the gap between the maximal weight of disagreement object pairs and the weight one of agreement pairs. When α approaches zero, the contribution of disagreement pairs to spARI becomes negligible, so spARI becomes close to ARI. In contrast, as α increases toward one, the accumulated weights of disagreement pairs become larger, resulting in a relatively large difference between spARI and ARI in this example. In practice, we set the default value of α to 0.8 to ensure a reasonable positive gap between the weights of disagreement and agreement pairs, which can also capture meaningful spatial information.

**Figure S2:**
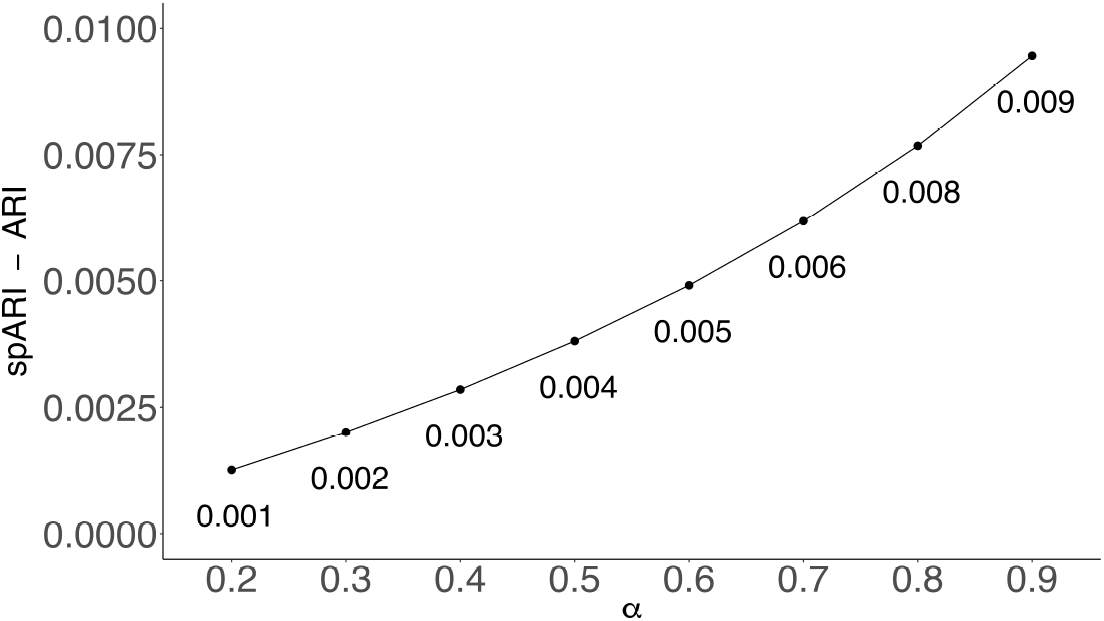
The differences between spARI and ARI of the first clustering result in the third simulation study in the main text, and the parameter α varies from 0.2 to 0.9 with gap 0.1.

## Web Appendix E: Analytical forms of spRI between two similar clusterings

Specifically, for any two groups *U*_*r*_ and 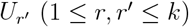 in the reference, we further assume that the limit of the arithmetic mean of *f*_*ij*_ or *h*_*ij*_ on *U*_*r*_ and 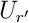 exist when the object size m per group increases. In other words,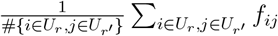 converges to µ_*f,U*_ (*r, r*^′^), and 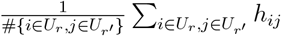 to µ_*h,U*_ (*r, r*^′^), as m goes to positive infinity.

In the first circumstance, we consider the following four operations on the reference partition **U** to generate four different clustering partitions **V**, respectively, as illustrated in Figure S3. For each operation, we demonstrate its specific spRI value between **U** and **V**, as well as its limits when either m or k tends to infinity in the rows (A1)-(A4) in Table S1.

(A1) Merge two groups *U*_*r*_ and 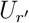 into a single cluster.

(A2) Split one group in **U** into two clusters *V*_*c*_ and 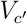 with each containing an equal number of objects (assuming m is even).

(A3) Split one group *U*_*r*_ into m clusters, each containing exactly one object.

(A4) Select one object from each group in **U**, and assign them together to a new cluster of size k. Let *U*(*i*) denote the original group in **U** to which object *i* belongs. Let ϕ_*V*_ be 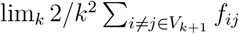, which is assumed to exist.

In the second circumstance, we apply the three following types of operations to the reference partition **U**, and each operation generates two different clustering results **V**_1_ and **V**_2_ on different parts of **U**, as illustrated in Figure S4. We then compute the spRI values between **V**_1_ and **V**_2_ as well as its limits in the rows (B1)-(B3) in Table S1.

(B1) Move an object a into two different reference groups, *U*_*s*_ and *U*_*t*_, respectively. Denote the clusters containing object a in **V**_1_ and **V**_2_ by *V*_1*c*_ and 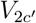, respectively.

**Figure S3:**
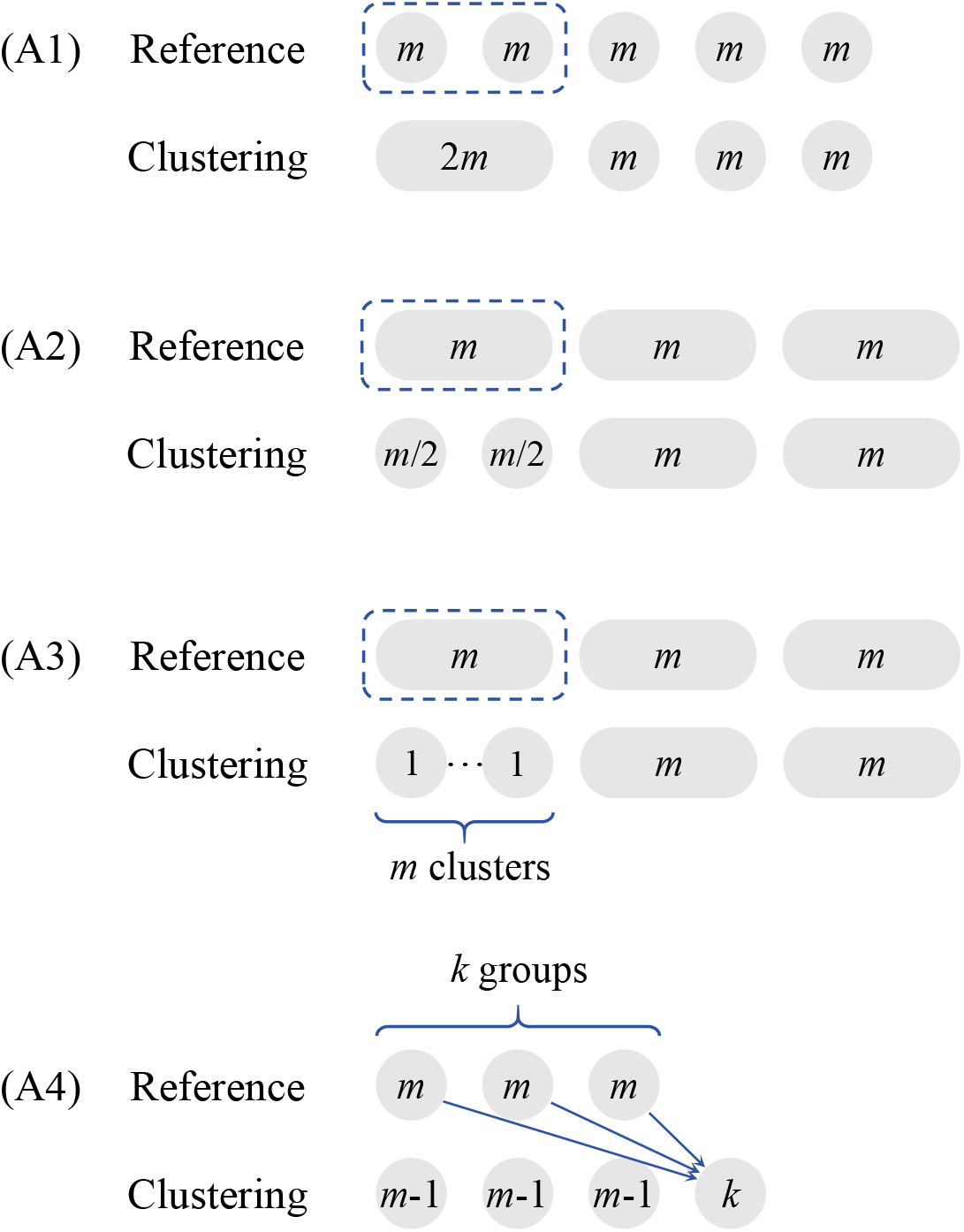
Illustrations of the four operations (A1)–(A4) applied to the reference partition, each resulting in a distinct clustering partition.

**Figure S4:**
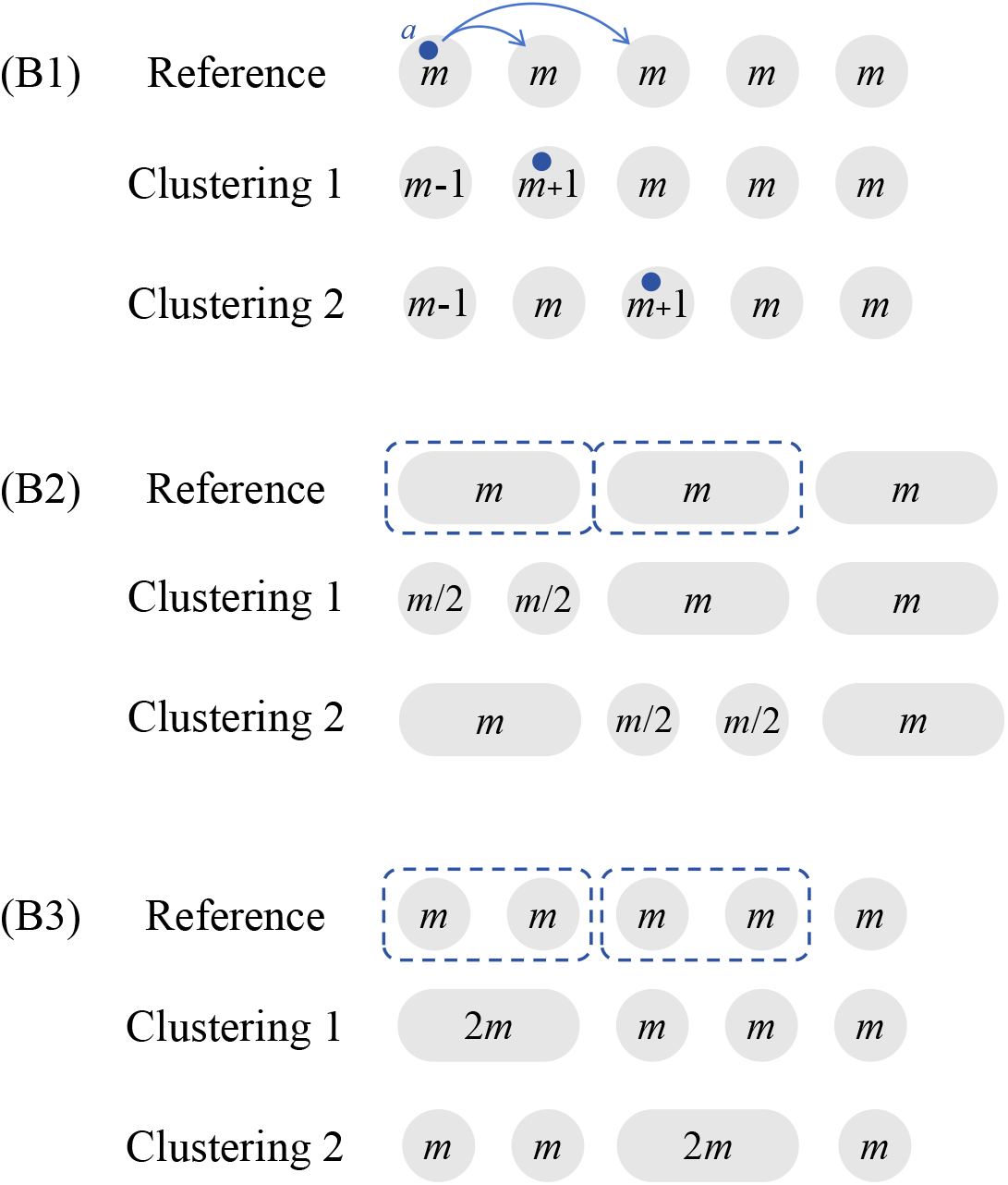
Illustrations of the three operations (B1)–(B3) applied to the reference partition, each resulting in two distinct clustering partitions.

**Figure S5:**
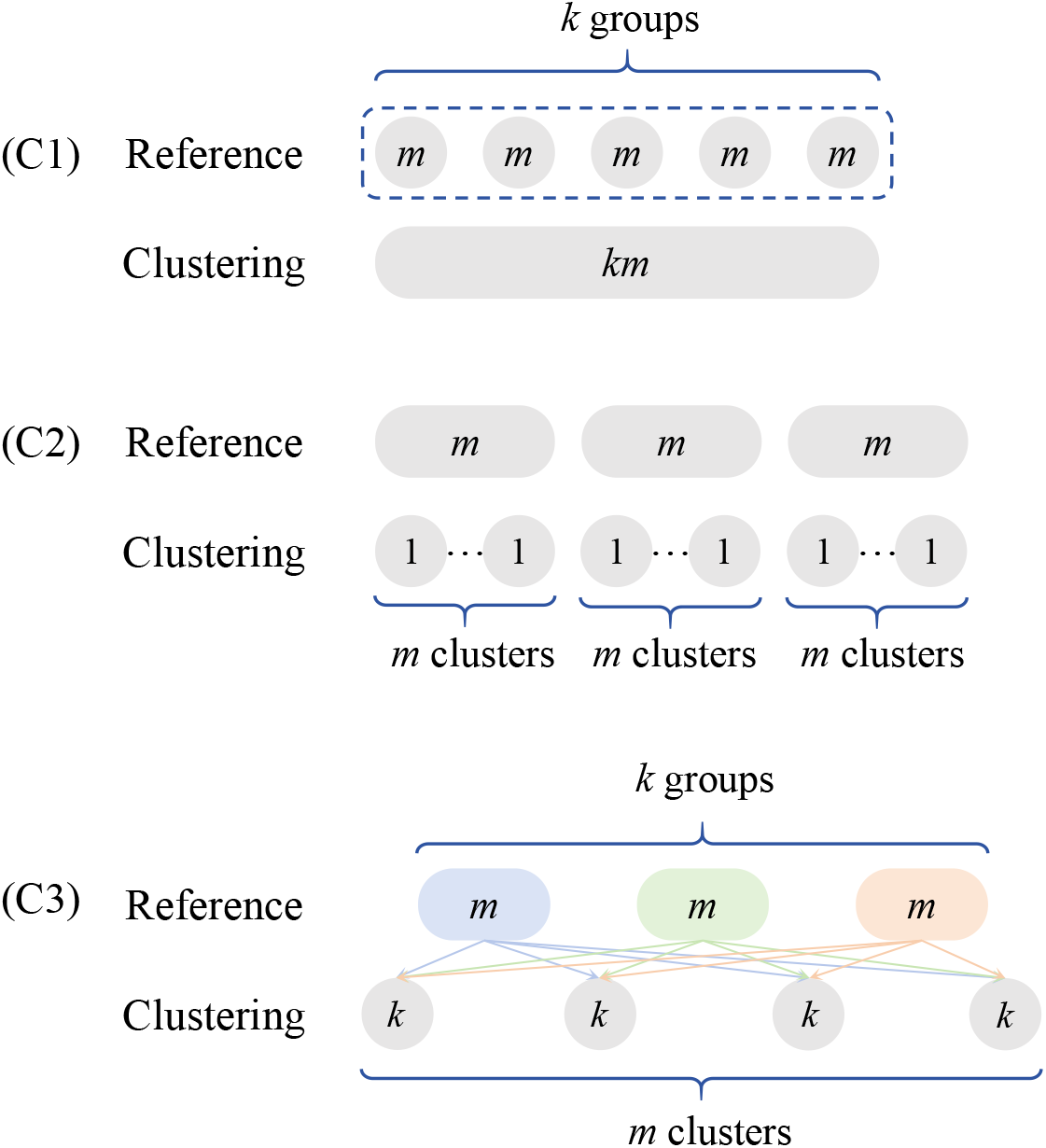
Illustrations of the three operations (C1)–(C3) applied to the reference partition, each resulting in a distinct clustering partition.

**Table S1:** Forms of spRI and its limits for operation cases (A1)-(A4), (B1)-(B3), and (C1)-(C3).

| Cases | spRI | $m \rightarrow +\infty$ | $k \rightarrow +\infty$ |
| --- | --- | --- | --- |
| (A1) | $\frac{k^2 - 2 - \frac{k}{m} + \frac{2}{m^2} \sum_{i \in U_r, j \in U_{r'}} f_{ij}}{k^2 - \frac{k}{m}}$ | $\frac{k^2 - 2 + 2\mu_{f,U}(r, r')}{k^2}$ | 1 |
| (A2) | $\frac{(2k^2 - 1) - \frac{2k}{m} + \frac{4}{m^2} \sum_{i \in V_c, j \in V_{c'}} h_{ij}}{2k^2 - \frac{2k}{m}}$ | $\frac{2k^2 - 1 + \mu_{h,V}(c, c')}{2k^2}$ | 1 |
| (A3) | $\frac{k^2 - 1 - \frac{k-1}{m} + \frac{2}{m^2} \sum_{i \neq j \in U_r} h_{ij}}{k^2 - \frac{k}{m}}$ | $\frac{k^2 - 1 + \mu_{h,U}(r, r)}{k^2}$ | 1 |
| (A4) | $\frac{m^2 - \frac{3m}{k} - 1 + \frac{3}{k} + \frac{2}{k^2} \sum_{i \neq j \in V_{k+1}} f_{ij} + \frac{2}{k^2} \sum_{i \in V_{k+1}, j \in U(i) \setminus \{i\}} h_{ij}}{m^2 - \frac{m}{k}}$ | 1 | $\frac{m^2 - 1 + \phi_V}{m^2}$ |
| (B1) | $\frac{k^2 m - k - 4 + \frac{2}{m} \sum_{j \in V_{2c'} \setminus \{a\}} f_{aj} + \frac{2}{m} \sum_{j \in V_{1c} \setminus \{a\}} h_{aj}}{k^2 m - k}$ | 1 | 1 |
| (B2) | $\frac{2k^2 - \frac{2k}{m} - 2 + \frac{4}{m^2} \sum_{i \in V_{1c}, j \in V_{1c'}} f_{ij} + \frac{4}{m^2} \sum_{i \in V_{2c}, j \in V_{2c'}} h_{ij}}{2k^2 - \frac{2k}{m}}$ | $\frac{2k^2 - 2 + \mu_{f,V_1}(c, c') + \mu_{h,V_2}(c, c')}{2k^2}$ | 1 |
| (B3) | $\frac{k^2 - \frac{k}{m} - 4 + \frac{2}{m^2} \sum_{i \in U_t, j \in U_{t'}} f_{ij} + \frac{2}{m^2} \sum_{i \in U_r, j \in U_{r'}} h_{ij}}{k^2 - \frac{k}{m}}$ | $\frac{k^2 - 4 + 2\mu_{f,U}(t, t') + 2\mu_{h,U}(r, r')}{k^2}$ | 1 |
| (C1) | $\frac{1 - \frac{1}{m} + \frac{2}{k} \sum_{r < r'} \frac{1}{m^2} \sum_{i \in U_r, j \in U_{r'}} f_{ij}}{k - \frac{1}{m}}$ | $\frac{1 + \frac{2}{k} \sum_{r < r'} \mu_{f,U}(r, r')}{k}$ | $\bar{\mu}_U$ |
| (C2) | $\frac{k^2 - k + \sum_r \frac{2}{m^2} \sum_{i \neq j \in U_r} h_{ij}}{k^2 - \frac{k}{m}}$ | $\frac{k^2 - k + \sum_r \mu_{h,U}(r, r)}{k^2}$ | 1 |
| (C3) | $\frac{(k-1)(m-1) + \frac{2}{km} [\sum_r \sum_{i \neq j \in U_r} h_{ij} + \sum_c \sum_{i \neq j \in V_c} f_{ij}]}{km - 1}$ | $\frac{k-1 + \frac{1}{k} \sum_r \mu_{h,U}(r, r)}{k}$ | $\frac{m-1 + \frac{1}{m} \sum_c \mu_{f,V}(c, c)}{m}$ |

(B2) Split different groups in **U** into two clusters of equal size (assuming m is even), respectively. Denote the two clusters in **V**_1_ and **V**_2_ by 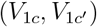 and 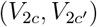, respectively.

(B3) Merge two distinct groups in **U**. Denote the two groups that are merged into a single cluster in **V**_1_ by *U*_*r*_ and 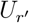, and those that are merged in **V**_2_ by *U*_*t*_ and 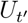.

In the third circumstance, we consider the following three operations on the reference partition that result in a clustering partition **V**, which are illustrated in Figure S5. For each operation, the values of spRI between **U** and **V** as well as its limits when either m or k tends to infinity are shown in the rows (C1)-(C3) in Table S1.

(C1) Merge all groups in the reference partition into a single cluster. We let

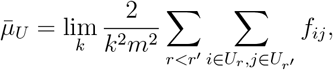

which is assumed to exist.

(C2) Split all groups in the reference partition into km clusters, each containing one object.

(C3) Reorganize all objects into m “opposite” clusters, each containing k objects that come from the k different groups in the reference partition.

## Web Appendix F: Calculations in the sensitivity of spRI

The spRI using the subsample *τ* Ω, denoted by spRI_*τ*_, is

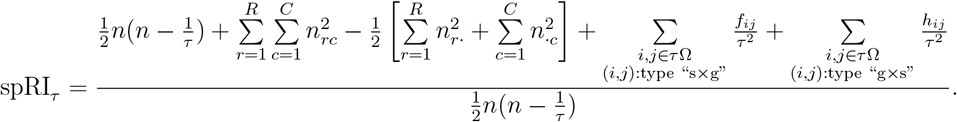

For the whole object set Ω, the spRI is

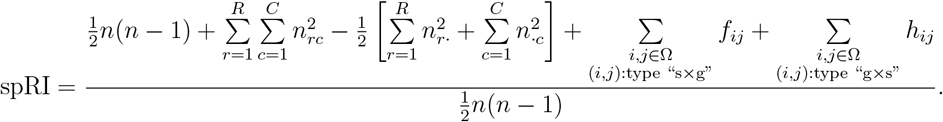

Subsequently, the difference in spRI between the subsample *τ* Ω and the whole sample Ω is given by

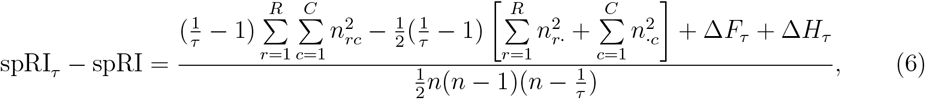

where

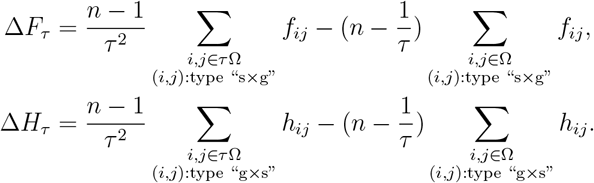

The spRI difference in Equation (6), spRI_*τ*_ − spRI, measures the sensitivity of spRI to the subsampling proportion *τ*. As expected, it approaches zero as the proportion *τ* increases to one. Let 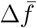 denote the difference between the mean value of *f*_*ij*_ over all type “s×g” pairs of objects in *τ* Ω and that in Ω, and 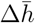 represents the difference between the mean value of *h*_*ij*_ over the type “g×s” pairs of objects in *τ* Ω and that in Ω. Recall that the type “s×g” object pair number and the type “g×s” object pair number in Ω are *N*_21_ and *N*_12_, respectively. According to Equations (1) and (2) in the main text, the corresponding numbers of such pairs in *τ* Ω are *τ* ^2^*N*_21_ and *τ* ^2^*N*_12_, respectively. The following proposition provides an upper bound on the spRI difference.

### Proposition S2.

*Assume that* 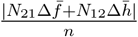 *converges to zero as n increases to infinity. Then the subsampling approximation error is given by* 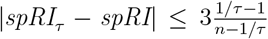 *when n is large enough*.

*Proof*. The Equation (6) can be rewritten as

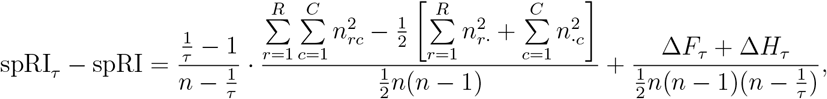

We denote the first term on the right hand side of this equation as T_1_ and the second term as T_2_. By the definition of RI, we have 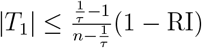. Next, we aim to derive an upper bound for T_2_. Define that

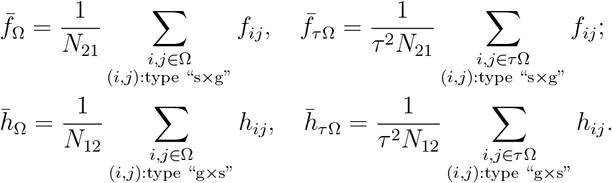

Since functions *f* and *h* take values in the interval [0, α], it is easy to verify that 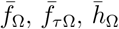 and 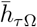 all lie in the range of [0, 1]. Then Δ*F*_*τ*_ and Δ*H*_*τ*_ can be reformulated as

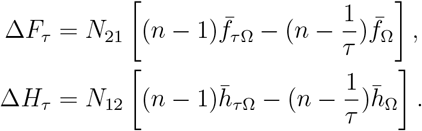

Recall that 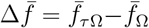 and 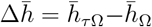. This gives us 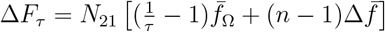 and 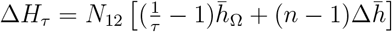. Substituting these values into *T*_2_, we obtain that

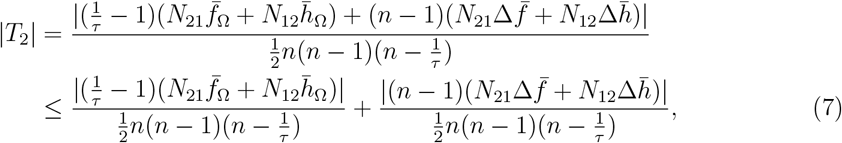

where the inequality follows from the triangle inequality. Denote the first and second term on the right hand side of Inequality (7) as |*T*_21_| and |*T*_22_|, respectively. For T_21_, since 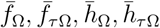 are bounded in the interval [0, 1], we have

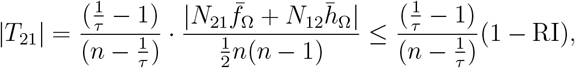

where the last inequality follows from the definition of RI. For T_22_, since we assume that 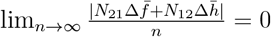, there exists a positive integer *n*_0_ such that when 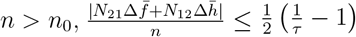. This implies that

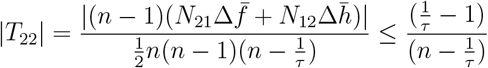

Substituting the results into Inequality (7), we have

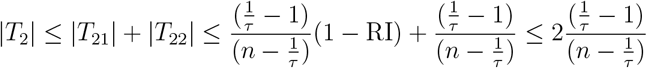

Finally, combining the upper bounds on |*T*_1_| and |*T*_2_|, we can obtain

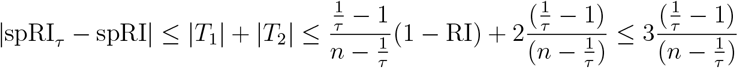

## Web Appendix G: Detailed steps in the permutation test

Given a significance level δ, the permutation testing procedure consists of the following four steps.

Step1 Randomly permute the reference labels and the clustering labels of all the objects, respectively. This ensures that the number of groups/clusters and the number of objects within each group/cluster remain the same as those in the original reference and clustering partitions. In other words, *R, C, n*_*r*·_, and *n*_·*c*_ do not change with the permutations.

Step2 Compute the simulated spARI value based on the permuted reference and clustering, denoted by spARI_sim_.

Step3 Repeat steps 1 and 2 *T* times to obtain *T* simulated spARI values.

Step4 Compute the proportion of simulated values satisfying |spARI_sim_ − ψ| ≥ |spARI_obs_ − ψ|. This proportion is taken as the *p*-value of this testing. If the *p*-value is smaller than the significance level δ, we have enough evidence to reject the null hypothesis, indicating that the observed spARI is significantly different from ψ.

The detailed steps in the algorithm are presented in Algorithm 1. Specifically, the notation spARI 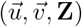 denotes the computation of spARI given the reference labels 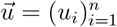, the clustering labels 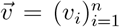, and the spatial coordinate matrix **Z**. The indicator function 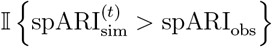 equals one when 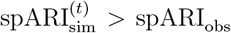 and equals zero otherwise.

We also added an R function “perm test” to automatically execute this permutation testing procedure.

Moreover, we incorporated an additional simulation case to provide some evidences that the proposed permutation test described above can control type I error well. In this scenario, the synthetic dataset consists of 200 objects, with the reference partition assigning them into two groups of sizes 1 and 199, respectively, and the clustering partition separating them into two clusters of sizes 90 and 110, respectively. The observed spARI value is close to zero (−0.0002), and the corresponding p-value derived from the R function “perm test” is 0.59, indicating that we do not have strong evidences to reject the null hypothesis spARI = 0 in the significance level δ = 0.1. Furthermore, we repeated the permutation test 100 times under the same setting, and the resulting p-values range from 0.43 to 0.64, none of which leads to an incorrect rejection of the null hypothesis. This demonstrates that the proposed hypothesis testing in Algorithm 1 effectively controls type I error.

### Algorithm 1: Permutation test for spARI

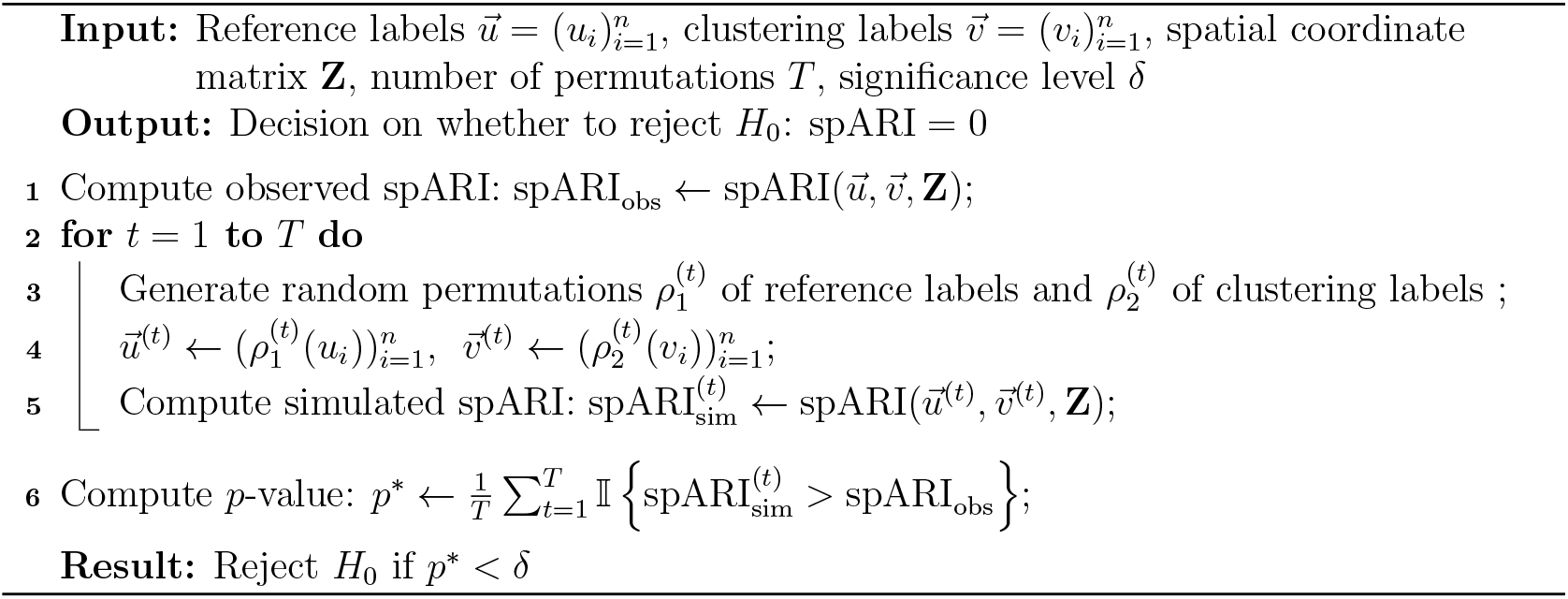

## Web Appendix H: Computational efficiency of spARI

We investigated the computational efficiency of spARI by varying the number of objects. Specifically, we generated ten simulated datasets with the object number *n* from 1,000 to 10,000 in increments of 1,000. In each dataset, all objects were partitioned into four reference groups, and half of them were set to be misclustered, which forms the clustering partition. Figures S6(a) and (b) demonstrate the boxplots of the computational time of spARI in seconds across ten replicates for each value of *n*. The numerical experiments were run on a MacBook Air powered by Apple M4 CPU with 16GB of RAM. We observe that in the setting where the spatial coordinates are provided, the runtime remains below one second for *n* ≤ 7, 000. If we use a precomputed distance matrix as input, the runtime stays under one second even for *n* ≤ 8, 000. Here, a distance matrix records the pairwise distances among all objects. Overall, employing a distance matrix can lead to a faster calculation compared to using spatial coordinates, which is expected as the computation of pairwise distances from coordinates becomes increasingly time-consuming when *n* is large. Accordingly, the R package of spARI supports the direct input of a distance matrix by users.

**Figure S6:**
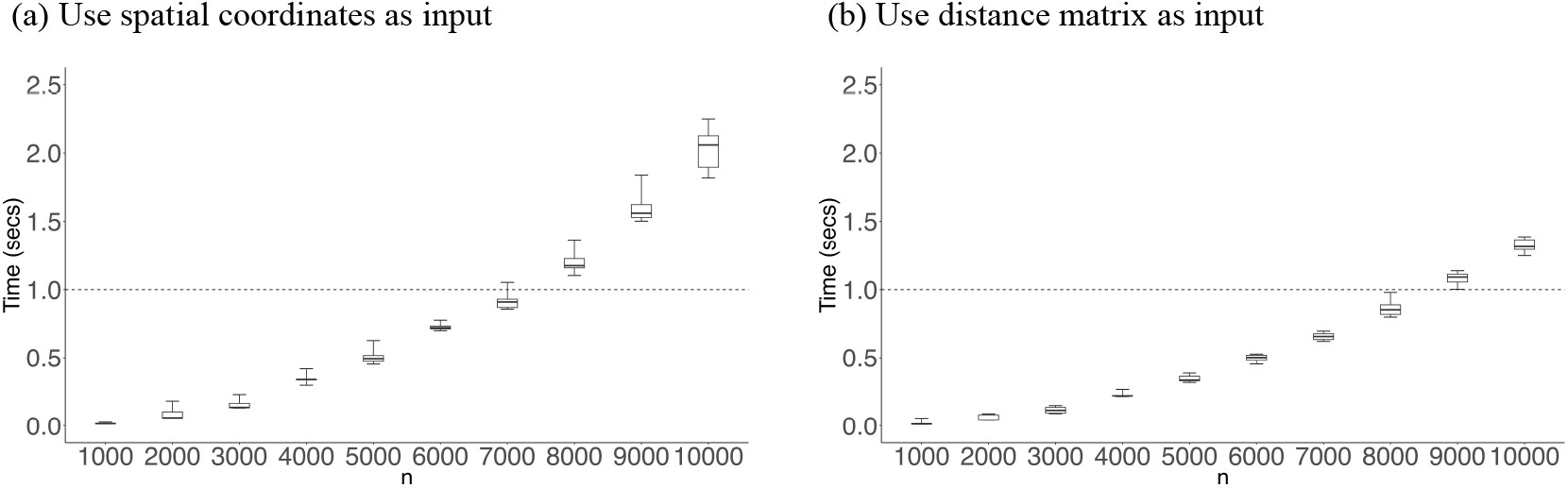
The boxplots of computational time of spARI in seconds for different object numbers when users provide (a) the spatial coordinates and (b) the distance matrix.

When *n* = 100, 000, the full dense distance matrix requires over 16GB of memory, making spARI incomputable in common laptops. To address this issue, we may store the distance matrix in a sparse format. We simulate a large-scale scenario by generating a synthetic ST dataset with *n* = 100, 000 cells. The spatial coordinates of these cells were uniformly sampled from the [0, 1] × [0, 1] region, and each cell was assigned to one of 15 distinct cell types. Due to the large size of the dataset, we focused only on the local spatial neighborhood of these cells. For example, we only computed the Euclidean distances for each cell’s five nearest neighbors. Since the distance matrix need to be symmetric, a distance between cells *i* and *j* was recorded in the sparse distance matrix if either *i* or *j* was among the other’s five nearest neighbors. The resulting sparse matrix contained approximately 600,000 nonzero entries. We then randomly selected 10,000 cells and manually assigned their cell type labels to construct one clustering partition. The reference partition, the clustering partition, and the sparse distance matrix was passed to the R function to compute the spARI value (with all other arguments set to default values). We repeated the computational process ten times, and the median runtime is 4.8 seconds. This shows that spARI is scalable to very large-scale ST data if users are able to provide the distance matrix in a sparse format.

## Web Appendix I: Biological interpretations for clustering results of BayesSpace and BANKSY

BayesSpace incorporates a Markov random field prior to induce spatial dependence among neighboring spots, which encourages them to be assigned to the same cluster. This may lead to integral spatial domains that are more consistent with the layered organization of the cortex. Maynard et al. (2021) reported that the marker gene *CCK* is highly expressed across the annotated Layers 2, 3 and 6, suggesting that these layers may share transcriptomic features or functional similarities. This observation may help explain why BayesSpace identified the cluster C4 that combines spots from these three layers. In contrast, BANKSY tends to produce smaller, spatially fragmented regions, as marked by the dashed circles in Figure 2(a) of the main text. From a biological perspective, such fragmentation may disrupt the regular layered architecture of the cortex in the ground truth. Therefore, the biological interpretations support our results that BayesSpace achieves a higher spARI value due to its better ability to capture spatial compactness than BANKSY.

In addition, we conducted permutation tests for the spARI values obtained from BayesSpace and BANKSY. As previously reported, their spARI values are 0.3594 and 0.3421, respectively. Both clustering results yielded p-values zero, providing strong evidences of significant associations with the reference partition. These results confirm that each method captures some meaningful spatial structures relative to the ground truth.

We also applied the hypothesis testing to the clustering results in the third simulation study (Section 4.3 in the main text). Recall that the spARI values of the two clustering results are 0.672 and 0.632. Implementing the permutation test based on Algorithm 1, both the clustering results obtained p-values of zero, strongly indicating their significant associations with the reference partition.

## Web Appendix J: Application of spARI to a geographical dataset

We use the social media geographical dataset in Kang et al. (2022) to demonstrate the application of spRI and spARI. This dataset contains 7,604 records, each with longitude and latitude geographical coordinates. The ground truth for position categories is shown in Figure S7(a) and serves as the reference partition, where the blue dots represent workplaces and the yellow dots denote home positions. Figure S7(b) demonstrates the clustering result estimated by Kang et al. (2022), along with its corresponding values of RI, spRI, ARI, and spARI.

**Figure S7:**
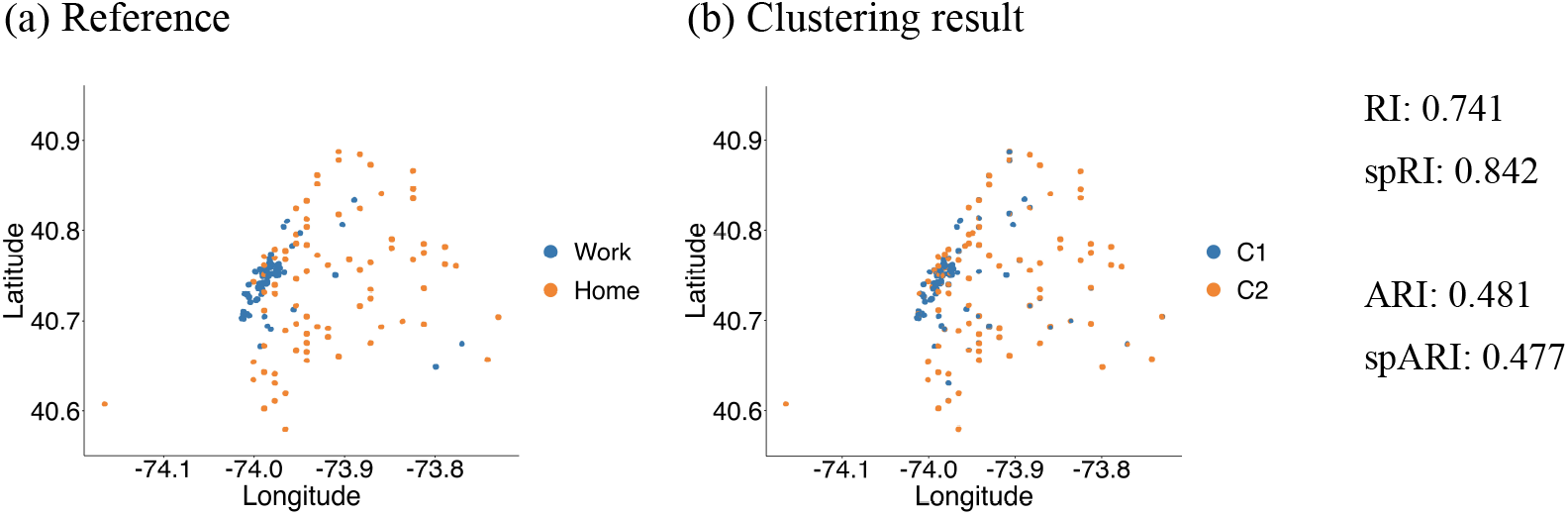
(a) The reference partition of the 7,604 social media records with geographical coordinates. (b) The estimated clustering partition. The values of RI, spRI, ARI, and spARI for the clustering result is shown at the right of panels.

Moreover, spARI is also compatible with spatial data characterized by an undirected graph structure, where each node corresponds to an object and edges denote connections. In this setting, the spatial proximity between any two objects is represented by an adjacency matrix, where the (*i, j*)-th entry equals one if objects *i* and *j* are connected and equals zero otherwise. This 0-1 symmetric adjacency matrix can be viewed as a special case of a distance matrix.

